# The GDF15-GFRAL pathway is dispensable for the effects of metformin on energy balance

**DOI:** 10.1101/2022.02.16.480373

**Authors:** Anders B. Klein, Trine S. Nicolaisen, Kornelia Johann, Andreas M. Fritzen, Cecilie V. Mathiesen, Cláudia Gil, Nanna S. Pilmark, Kristian Karstoft, Martin B. Blond, Jonas S. Quist, Randy J. Seeley, Kristine Færch, Jens Lund, Maximilian Kleinert, Christoffer Clemmensen

## Abstract

Metformin is a blood glucose lowering medication with physiological effects that extend beyond its anti-diabetic indication. Recently, it was reported that metformin lowers body weight via induction of growth differentiation factor 15 (GDF15), which suppresses food intake by binding to the GDNF family receptor α-like (GFRAL) in the hindbrain. At the same time, we demonstrated that recombinant GDF15 suppresses voluntary exercise in a GFRAL-dependent fashion. Here, we corroborate that metformin increases circulating GDF15 in mice and humans, but that it does not reduce voluntary running activity in mice. Unexpectedly, we fail to confirm previous reports that the GDF15-GFRAL pathway is necessary for the weight-lowering effects of metformin. Instead, our studies in wild-type, GDF15 knockout and GFRAL knockout mice suggest that the GDF15-GFRAL pathway is dispensable for the effects of metformin on energy balance. The data presented here question whether metformin is a sufficiently strong stimulator of GDF15 to drive anorexia and weight loss and emphasize that additional work is needed to untangle the relationship among metformin, GDF15 and energy balance.

## INTRODUCTION

Growth differentiation factor 15 (GDF15), a cellular stress signal that is expressed and secreted from various tissues, has caught recent attention as a promising target for treatment of obesity. Recombinant GDF15 reduces food intake via the GDNF family receptor α-like (GFRAL) receptor located in the area postrema (AP) and the nucleus tractus solitaris (NTS) of the hindbrain (Hsu et al., 2017; Mullican et al., 2017; Xiong et al., 2017; Yang et al., 2017). The food intake lowering effects of GDF15 extent to mice, rodents, and nonhuman primates (Hsu et al., 2017; Mullican et al., 2017; Xiong et al., 2017; Yang et al., 2017). In 2017, GDF15 emerged as a top biomarker candidate for the widely prescribed anti-diabetic medication metformin (Gerstein et al., 2017). Metformin moderately lowers body weight in humans (Diabetes Prevention Program Research, 2012) and this is suggested to be a key effect for the glycemic benefits (Lachin et al., 2007). Recently, it was reported that metformin increases circulating GDF15 levels in mice and humans and that the weight-lowering effects of metformin depend on functional GDF15-GFRAL signaling in mice (Coll et al., 2020; Day et al., 2019). At the same time, we demonstrated that recombinant GDF15 suppresses voluntary exercise in a GFRAL-dependent fashion (Klein et al., 2021). Given that metformin increases perceived exertion during exercise in humans (Das et al., 2018), we wanted to understand whether this phenomenon is related to GDF15.

## RESULTS

### Metformin increases circulating GDF15 in a diet-dependent manner

To study GDF15-GFRAL-dependent effects of metformin on exercise behavior and appetite, we orally administered vehicle or metformin (400 mg/kg) in lean chow-fed female wild type (WT) and GFRAL knockout (KO) mice (Figure 1A). In line with previous results (Coll et al., 2020; Day et al., 2019), metformin increased plasma GDF15 by 2-fold in both genotypes (Figure 1B). Despite elevating circulating GDF15, metformin did not influence running distance over 24 hours in WT or in GFRAL KO mice (Figure 1C). This finding contrasts reports suggesting that metformin enhances the perception of exertion during exercise (Das et al., 2018; Kristensen et al., 2019; Pilmark et al., 2021c). We did not observe decreased chow intake in metformin-treated mice (Figure 1D), suggesting that exercise might mask the effects of metformin to lower appetite. Previous studies have reported that in high-fat diet (HFD)-fed mice metformin treatment promotes a more pronounced induction in plasma GDF15, but it is unknown whether this relates to i) changes in nutrient intake (e.g., higher fat intake), ii) a positive energy balance or iii) increased adiposity that manifests with long-term HFD intake. To determine this, we measured plasma levels of GDF15 3 hours after a single oral administration of metformin (400 mg/kg) in male mice on chow diet and in male mice switched from chow to HFD diet for 1 day, 3 days, 10 days or 21 days (Figure 1E, Figure S1A). Metformin doubled the plasma level of GDF15 in mice on chow diet (Figure 1F), but already after 3 days on HFD diet, metformin induced 2.7-fold higher plasma GDF15 levels compared to the chow-fed condition (Figure 1F). Both the liver and the gastrointestinal tract are targeted by metformin (Coll et al., 2020; Yang et al., 2021), and these two tissues might be primary sources of circulating GDF15 in metformin-treated mice. In contrast to findings in primary mouse hepatocytes (Coll et al., 2020; Day et al., 2019), oral metformin administration decreased *Gdf15* expression in the liver of both chow-fed mice and mice exposed to HFD for 1 and 3 days (Figure 1G). Instead, metformin increased *Gdf15* expression in duodenum and colon of both chow-fed mice and HFD-fed mice (Figure 1H-I). In the kidney, metformin had no effect on *Gdf15* expression in chow-fed mice, but it increased *Gdf15* expression by 3-fold after 3 days on HFD (Figure 1J). These observations emphasize the importance of diet composition for the effects of metformin on GDF15 secretion. They also indicate that in mice the potentiated metformin-induced release of GDF15 after 3 days on HFD originates from kidney. Given these insights, that the GDF15-response to metformin is largely dependent upon diet type (Figure 1K), we evaluated whether a greater GDF15-response to metformin in HFD-fed mice suppresses voluntary running. We switched female mice from chow to HFD and administered a single oral dose of metformin (300 mg/kg) five days later. Metformin resulted in a higher induction in plasma GDF15 in mice switched to HFD compared to mice kept on chow (Figure 1B versus Figure S1C). However, despite the potentiated GDF15 response, metformin did not lower voluntary running distance in HFD-fed mice (Figure S1D). Unexpectedly, metformin lowered food intake in exercising HFD-fed female mice independent of GFRAL (Figure S1E), suggesting that increased physical activity modulates the mechanism by which metformin lowers food intake.

**Figure 1.**
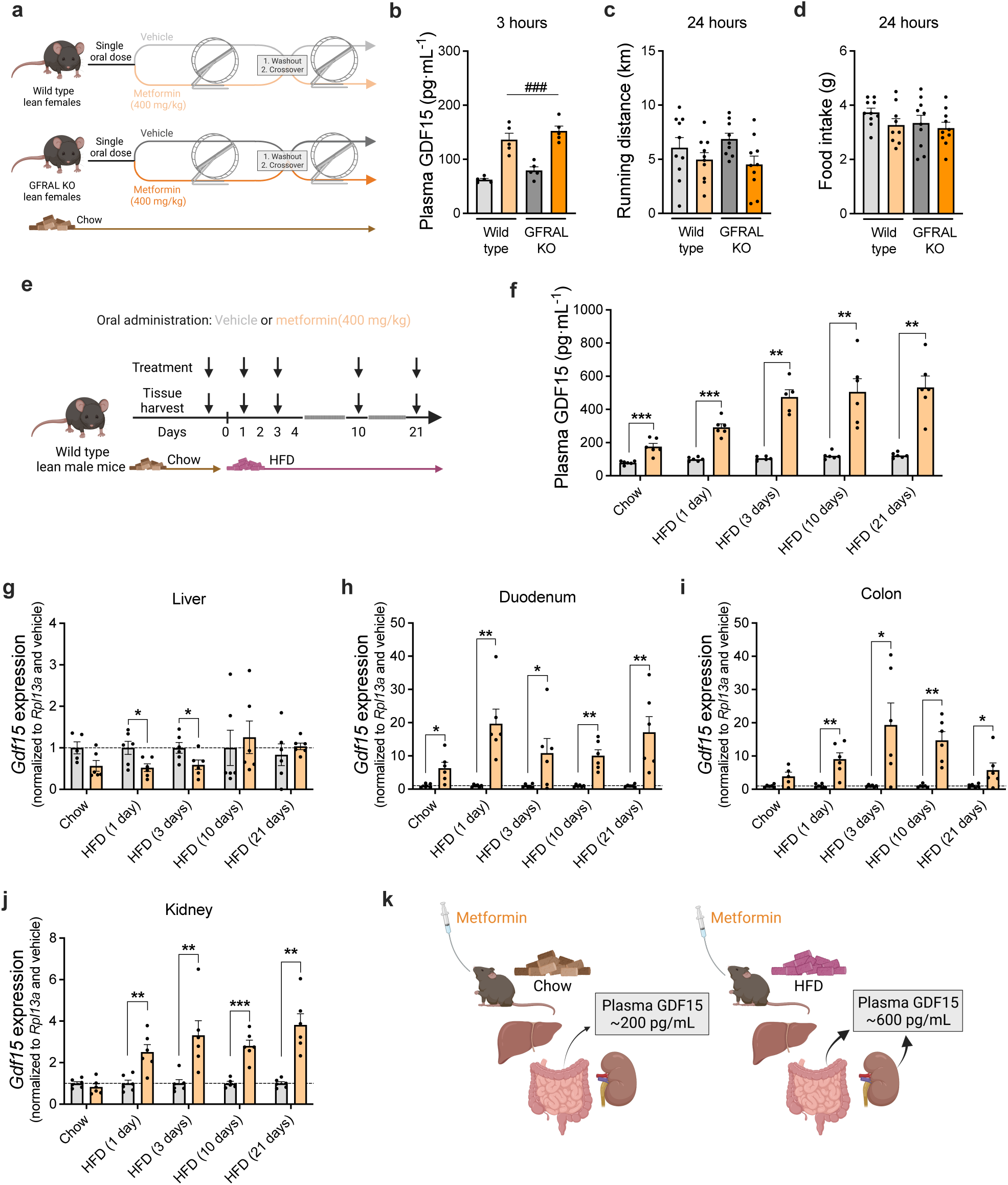
Metformin stimulates GDF15 tissue expression and circulating levels in a diet-dependent fashion. **a**, Illustration of study in WT and GFRAL KO female mice with access to running wheels. Mice were treated by oral gavage with metformin (400 mg/kg) or vehicle and running distance was measured over 24 hours. The experiment was repeated following an 11-day washout period, with vehicle mice receiving metformin and vice versa. **b**, Blood plasma GDF15 levels measured 3 hours following oral gavage with metformin (400 mg/kg) or vehicle. Data were analyzed by two-way ANOVA, ^###^p<0.001; main effect of metformin. **c**, Running distance after 24 hours. Data were analyzed by two-way RM ANOVA. **d**, Food intake after 24 hours following a single dose of metformin (400 mg/kg) or vehicle by oral gavage. Data were analyzed by two-way RM ANOVA. **e**, Illustration of study in which wild type male mice were fed chow or high-fat diet (HFD) for either 1, 3, 10, or 21 days and treated with either vehicle or metformin (400 mg/kg). **f**, Blood plasma GDF15 levels at designated time points after oral gavage of either vehicle or metformin. Data were analyzed by an unpaired two-tailed t-test, Mann-Whitney test or Welch’s test, depending on Gaussian distribution or equal variance, for each time point, **p<0.01, ***p<0.001. **g-j**, *Gdf15* expression in liver (**g**), duodenum (**h**) colon (**i**) and kidney (**j**) at different time-points. Data were analyzed by an unpaired two-tailed t-test, Mann-Whitney test or Welch’s test, depending on Gaussian distribution or equal variance, for each time point, *p<0.05, **p<0.01, ***p<0.001. **k**, Illustration of tissue contributions to blood plasma GDF15 in mice on either chow or HFD diet and treated with metformin by oral gavage.

### Metformin lowers body weight independent of the GDF15-GFRAL pathway

Intrigued by the observation that metformin induces anorexia in exercising GFRAL null mice on a HFD, we first wanted to ensure that we could replicate the necessity of the GDF15-GFRAL pathway for the weight lowering effect of metformin, using a published protocol (Coll et al., 2020). Accordingly, lean male WT and GFRAL KO mice (Figure 2A-D) or WT and GDF15 KO mice (Figure 2E-H) mice were switched from chow to a HFD diet for 3 days before receiving daily oral administration of metformin (300 mg/kg) or vehicle for 11 days (Figure 2A,E). Similar to previous findings (Coll et al., 2020; Day et al., 2019), this treatment elicited a ∼2-fold increase in circulating GDF15 3 hours after the last metformin treatment (Figure 2B,F), but in contrast to Coll et al. (Coll et al., 2020) metformin failed to lower body weight or to reduce food intake (Figure 2A-H) in both WT and transgenic mice. Importantly, and in addition to the induction of GDF15, metformin treatment lowered blood glucose in WT and GDF15 KO mice (Figure S2A). It is possible that differences in environment and/or experimental conditions (Kleinert et al., 2018; von Herrath et al., 2019) are responsible for the perplexing absence of the weight lowering effects of metformin compared to previous work. The difference might also relate to the use of relatively lean mice, in which the “window” for healthy weight loss is smaller. To test this, we studied the effects of metformin in diet-induced obese (DIO) mice. We first performed a six-day long dose-titration study (Figure 2I) showing that daily treatment of obese mice with metformin elicits a 3-4-fold increase in plasma GDF15 levels 3 hours following the last administration (Figure 2J). Concentration of GDF15 reached ∼1000 pg/mL in mice treated with daily oral doses of 300 mg/kg and 400 mg/kg metformin and, importantly, coincided with reduced body weight (Figure 2L), decreased food intake (Figure 2L) and improved glycemic control (Figure S2B). To test whether the GDF15-GFRAL pathway mediates these effects in obese mice, we gavaged DIO WT and DIO GFRAL KO mice as well as DIO WT and DIO GDF15 KO mice daily for seven days with 400 mg/kg metformin (Figure 2M and Figure 2Q). As expected, the treatment increased circulating GDF15 (Figure 2N and 2R) but surprisingly reduced body weight and food intake to the same extent in both WT and KO mice (Fig 2O-P and 2S-T). Since these findings contrast results from two previous studies (Coll et al., 2020; Day et al., 2019), we generated another cohort of DIO WT and GFRAL KO mice and performed a complementary acute experiment in which metformin was administered as a single oral dose (400 mg/kg) and 24 h food intake and weight change were determined (Figure S2C). Similarly, under these treatment conditions metformin increased plasma GDF15 and decreased food intake and body weight in both WT and GFRAL KO mice. (Figure S2D-F). In another additional cohort of DIO WT and DIO GDF15 KO mice, we tested whether metformin administered at the beginning of the light phase lowers body weight via GDF15 (Figure S2G). However, similar to the other studies in which metformin was given at the end of the light phase (Figure 2A,E,I,M,Q), the earlier administration of metformin lowered body weight, food intake and blood glucose to a similar extent in DIO WT and DIO GDF15 KO animals (Figure S2G-K). Collectively, our data suggest that GDF15-GFRAL is dispensable for the effects of metformin on food intake and body weight.

**Figure 2.**
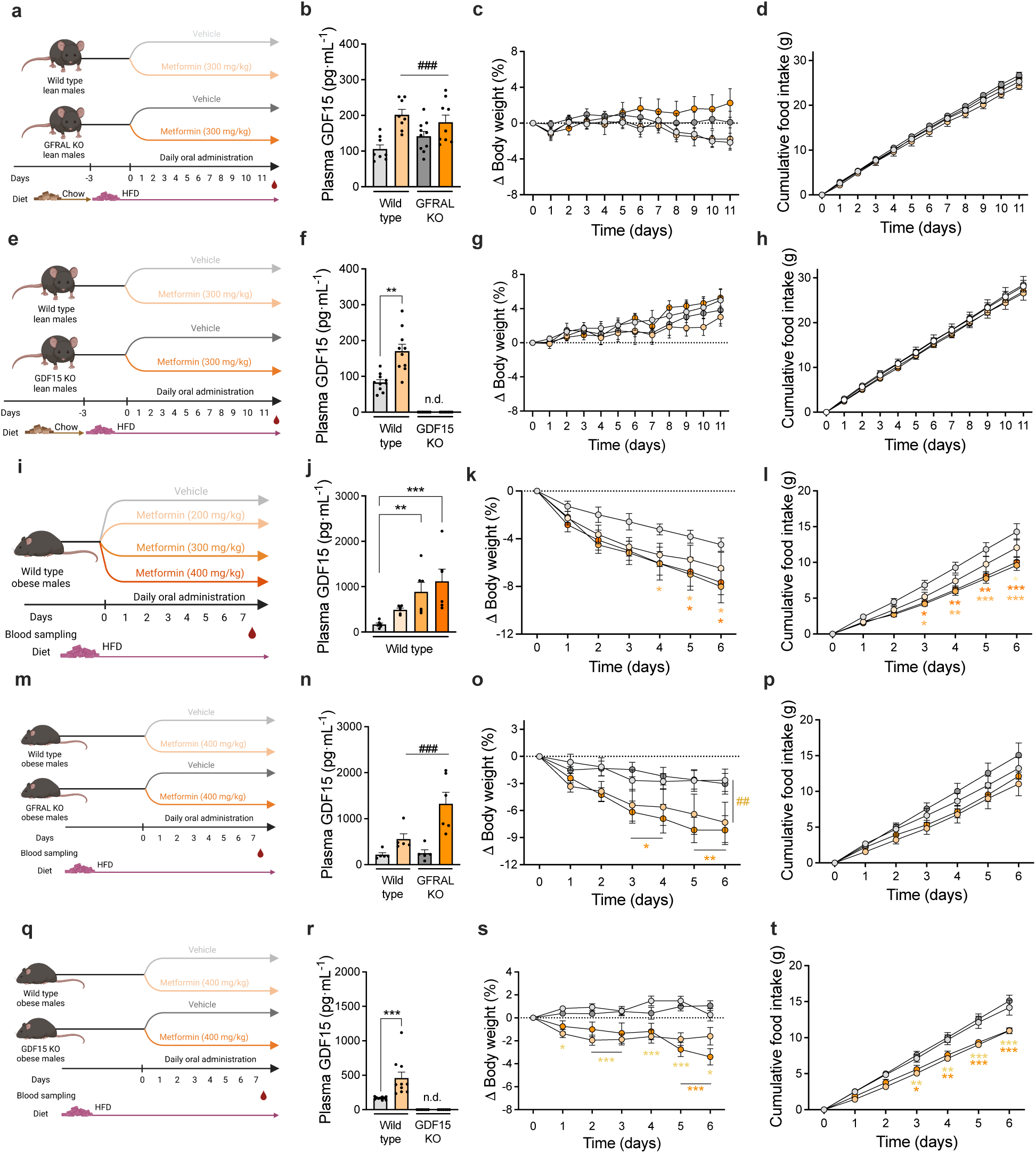
Metformin lowers body weight independently of the GDF15-GFRAL pathway in diet-induced obese mice. **a**, Illustration of study in male wild type and GFRAL knockout (KO) mice that were switched from chow diet to high-fat diet (HFD). Following 3 days on HFD, mice were treated daily by oral gavage for 11 days with metformin (300 mg/kg) or vehicle. **b**, Blood plasma GDF15 levels 3 hours following the final dose of vehicle or metformin. Data were analyzed by a two-way ANOVA, ^###^p<0.001; main effect of metformin. **c**, Change in body weight. Data were analyzed by a two-way RM ANOVA. **d**, Cumulative food intake. Data were analyzed by a two-way RM ANOVA. **e**, Illustration of study in male wild type and GDF15 knockout (KO) mice that were switched from chow diet to high-fat diet (HFD). Following 3 days on HFD, mice were treated by oral gavage for 11 days with metformin (300 mg/kg) or vehicle. **f**, Blood plasma GDF15 levels 3 hours following the final dose of metformin. The GDF15 plasma levels were below the detection limit for GDF15 KO mice (n.d. = not detectable). Data were analyzed by a Welch’s test within the wild type group, **p<0.01. **g**, Change in body weight. Data were analyzed by a two-way RM ANOVA. **h**, Cumulative food intake. Data were analyzed by a two-way RM ANOVA. **i**, Illustration of study in high-fat diet (HFD)-induced obese WT mice treated daily by oral gavage with metformin (200, 300 or 400 mg/kg) or vehicle for 7 days. **j**. Blood plasma GDF15 levels 3 hours following the final dose of metformin on day 7. Data were analyzed by a Kruskal-Wallis on Ranks test with Dunn’s multiple comparison test, **p<0.01, ***p<0.001. **k**, Change in body weight. Data were analyzed by a two-way RM ANOVA with Bonferroni multiple comparison test, *p<0.05. **l**. Cumulative food intake. Data were analyzed by a two-way RM ANOVA with Bonferroni multiple comparison test, *p<0.05, **p<0.01, ***p<0.001. **m**, Illustration of study in high-fat diet (HFD)-induced obese wild type and GFRAL knockout (KO) male mice treated by daily oral gavage with metformin (400 mg/kg) or vehicle for 7 days. **n**, Blood plasma GDF15 levels 3 hours following the final dose of metformin on day 7. Data were analyzed by a two-way ANOVA, ^###^p<0.001; main effect of metformin. **o**, Change in body weight. Data shown were analyzed by a two-way RM ANOVA with Bonferroni multiple comparison test, ^###^p<0.001; main effect of metformin within WT. *p<0.05, **p<0.01; effect of metformin within GFRAL KO. **p**, Cumulative food intake. Data were analyzed by a two-way RM ANOVA. **q**, Illustration of study in high-fat diet (HFD)-induced obese wild type and GDF15 knockout (KO) male mice treated by daily oral gavage of metformin (400 mg/kg) or vehicle for 7 days. **r**, Blood plasma GDF15 levels 3 hours following the final dose of metformin on day 7. The GDF15 plasma levels were below the detection limit for GDF15 KO mice (n.d. = not detectable). Data were analyzed by a Mann-Whitney test within wild type group, ***p<0.001. **s**, Change in body weight. Data were analyzed by a two-way RM ANOVA with Bonferroni multiple comparison test, *p<0.05, **p<0.01, ***p<0.001; effect of metformin within genotype. **t**, Cumulative food intake. Data were analyzed by a two-way RM ANOVA with Bonferroni multiple comparison test, *p<0.05, **p<0.01, ***p<0.001; effect of metformin within genotype.

### Metformin-induced GDF15 is not associated with body weight changes in individuals with overweight and prediabetes

Having observed that GDF15 is not universally required for metformin to reduce food intake in mice, we revisited the relationship of metformin, GDF15 and weight loss in humans. Individuals with overweight and prediabetes treated twice daily with 850 mg metformin for 13-weeks had GDF15 plasma levels that were 50% higher compared to a non-treated control group (Figure 3A). Despite this GDF15 increase, metformin was not associated with lower body weight compared with placebo (Figure 3B). We also failed to detect a previously reported^11,12^ association between changes in circulating GDF15 levels and changes in body weight (Figure 3C). Moreover, subjects with overweight/obesity (BMI 35.5±4.4 kg/m^2^) and prediabetes exposed to an exercise lifestyle intervention and metformin treatment (1000 mg twice daily) for 12 weeks (Pilmark et al., 2021a) displayed a ∼2-fold increase in GDF15 plasma levels (Figure S3A), while change in body weight was similar to the exercise control group (Figure S3B). No association between changes in plasma GDF15 levels and changes in body weight was detected in the study (Figure S3C). In contrast to previous findings, the clinical data presented here, suggest that metformin-induced increase in plasma GDF15 can be uncoupled from weight-lowering effects of metformin. Future studies with larger samples sizes are needed to clarify the putative association (and causation) between metformin, GDF15 and energy balance in humans. It may be that certain individuals or patient subgroups have an amplified responsiveness to the weight-lowering effects of metformin-induced GDF15.

**Figure 3.**
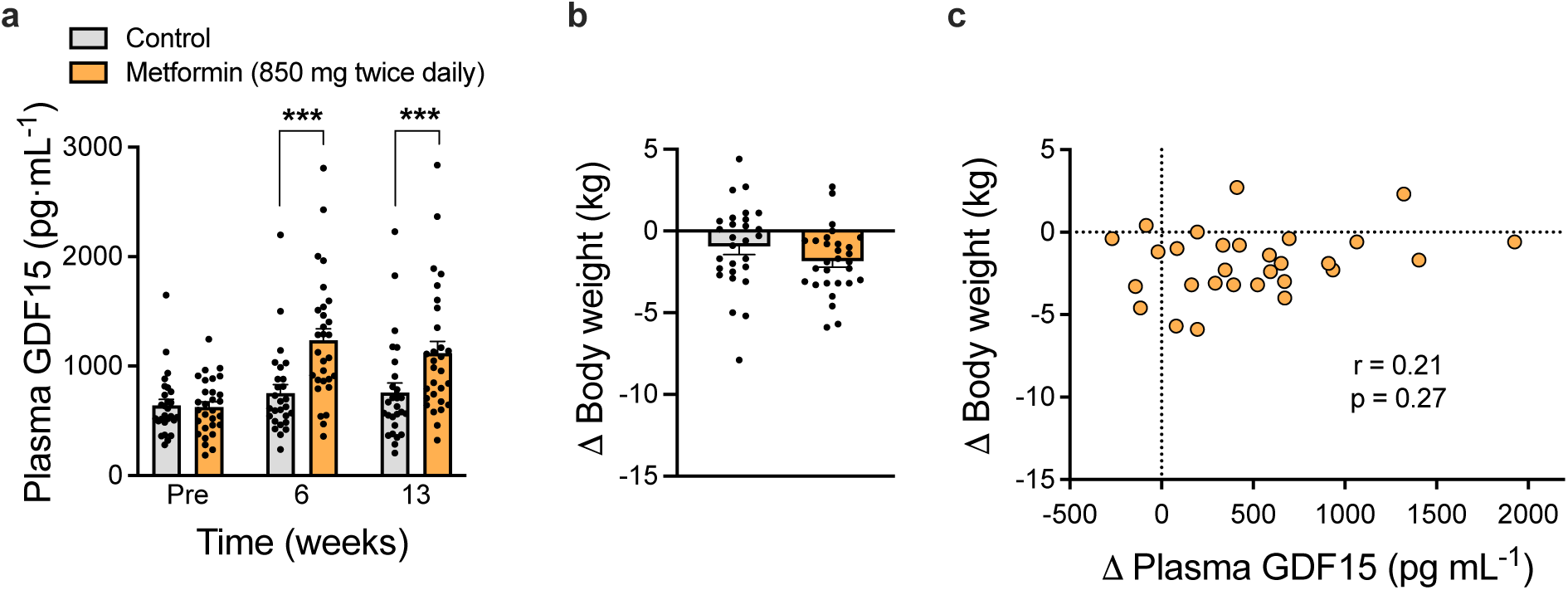
Metformin increases plasma GDF15 but is not associated with body weight loss in individuals with overweight and prediabetes. **a**, Plasma concentrations of GDF15 in individuals with overweight and prediabetes before and after 6- and 13-weeks of treatment with metformin or no treatment. Data were analyzed by a baseline constrained repeated measures regression model with plasma GDF15 as a function of age, sex, time and a group-by-time interaction (***p<0.001). **b**, Change in body weight following 13 weeks of treatment of metformin or no treatment. Data were analyzed by an unpaired two-tailed t-test between placebo and metformin **c**, Change in body weight versus change in plasma GDF15 levels in the subjects treated with metformin for 13 weeks. Data were analyzed by a Spearman correlation test.

## DISCUSSION

Here, we demonstrate that metformin does not reduce voluntary running behavior in mice. Moreover, and in contrast to two previous independent studies (Coll et al., 2020; Day et al., 2019), we find that metformin fails to lower body weight or prevent weight gain in lean HFD-fed mice. In diet-induced obese mice, metformin does lower body weight, however the GDF15-GFRAL pathway is dispensable for these effects. The mouse work was conducted in two transgenic loss-of-function models for the GDF15-GFRAL pathway (GFRAL KO and GDF15 KO) and at two different institutions/animal facilities. We also fail to confirm a previously reported association between metformin-induced GDF15 and weight loss in patients with overweight/obesity and prediabetes.

Drugs and chemicals that elicit sufficiently high levels of circulating GDF15 *can* drive weight loss in rodents by activating GFRAL-signaling in the hindbrain (Hsu et al., 2017; Sabatini et al., 2021; Worth et al., 2020). However, given the pleiotropic toxic nature of many potent GDF15 secretagogues, the direct contribution from elevated plasma GDF15 to the drug-induced anorectic response can be challenging to discern. For example, whereas weight loss following treatment with the chemotherapeutic drug, cisplatin, is ascribed to be partially driven by an increase in circulating GDF15 (Breen et al., 2020; Hsu et al., 2017), weight loss induced by endotoxin is not attenuated in GDF15-GFRAL loss-of-function mice, despite that endotoxin increases plasma GDF15 by a striking 20-50-fold in mice (Breen et al., 2021; Luan et al., 2019; Patel et al., 2021).

Collectively, the data presented here challenge recent findings that the GDF15-GFRAL pathway mediates the weight-lowering benefits of metformin in mice. Given that even subtle differences in environmental or experimental conditions influence study outcomes in rodents (Kleinert et al., 2018), it remains possible that under certain circumstances the GDF15-GFRAL pathway is required for the weight lowering effect of metformin. Yet, we show that this is not a robust universal effect, which is further underscored by the lack of association between metformin-induced GDF15 and weight loss in two human intervention studies. We encourage further studies focusing on GDF15-dependent and GDF15-independent effects to advance our understanding of the metabolic actions of metformin on mammalian energy balance. We also encourage studies focusing on identifying co-factors or pathways that may interact-or converge with the GDF15-GFRAL pathway to promote anorexia and weight loss.

## Supporting information

Combined Supplemental Data

## ACKNOWLEDGEMENTS

The authors thank Charlotte Svendsen for technical assistance. The authors thank the participants in the human studies. Human Study 1 (The PRE-D Trial) was funded by the Novo Nordisk Foundation, AstraZeneca AB, the Danish Innovation Foundation, the University of Copenhagen and Ascensia Diabetes Care Denmark ApS. This work was supported by research grants from Independent Research Fund Denmark (0134-00254B), the Lundbeck Foundation (Fellowship R238-2016-2859), the Deutsche Forschungsgemeinschaft (KL 3285/2-1), the German Center for Diabetes Research (DZD 82DZD03D03 & 82DZD03D1Y), and the Novo Nordisk Foundation (grant numbers NNF17OC0026114 & NNF19OC0055192). R.J.S. acknowledges support from: P01DK117821, DK119188, and P30DK089503. Novo Nordisk Foundation Center for Basic Metabolic Research is an independent Research Center, based at the University of Copenhagen, Denmark, and partially funded by an unconditional donation from the Novo Nordisk Foundation (www.cbmr.ku.dk) (Grant number NNF18CC0034900). Cartoon illustrations were created using BioRender.com.

## AUTHOR CONTRIBUTIONS

A.B.K. and CC conceptualized the project. A.B.K., T.S.N., K.J., A.M.F., C.V.M., C.R.E.G., N.S.P., K.K., M.B.B., J.S.Q., R.J.S., K.F., J.L., M.K., and C.C performed experiments and/or analyzed/interpreted data. A.B.K., K.J., J.L., M.K., and C.C., wrote the manuscript. All authors edited the manuscripts and provided comments.

## METHODS

### Human Study 1

Plasma samples were obtained from controls and metformin treated participants in the PRE-D Trial, described in detail elsewhere(Færch et al., 2021). All participants had a BMI >25kg/m^2^ and prediabetes defined by HbA1c between 39-47 mmol/mol. The trial was an investigator-initiated, randomized, controlled, open-label, four-arm (1:1:1:1), superiority trial performed at Steno Diabetes Center Copenhagen, Gentofte, Denmark from 2016 to 2019. The trial protocol was approved by the Ethics Committee of the Capital Region (H-15011398) and the Danish Medicines Agency (EudraCT: 2015-001552-30) and registered at ClinicalTrials.gov (NCT02695810). Approval of data storage was obtained from the Danish Data Protection Board (2012-58-0004). The trial was conducted in accordance with the Helsinki II declaration and Good Clinical Practice.

### Human Study 2

Plasma samples were obtained from a parallel-group, randomized clinical trial (Pilmark et al., 2021a). In brief, participants were randomly allocated to placebo + exercise training [PLA] or metformin + exercise training [MET]). Both investigators and participants were blinded to the treatment. All data were collected at Trygfondens Centre for Physical Activity Research at Rigshospitalet, Copenhagen (Pilmark et al., 2021b). The primary outcome was change in postprandial glucose, measured by mean glucose concentration during a 4-hour mixed meal tolerance test (MMTT). For further details, see ClinicalTrials.gov (NCT03316690).

### Animals

Wild type (WT) C57BL/6J male and female mice were obtained from commercial breeders (Janvier, FR). The GFRAL knockout (KO) and WT littermates were generated as previously reported (Frikke-Schmidt et al., 2019; Klein et al., 2021). One week before experimental treatment, mice were acclimatized to single housing. All experiments were done at 22°C with a 12:12 hour light-dark cycle. Mice had ad libitum access to water and chow diet (Altromin 1324, Brogaarden, Denmark), or when indicated a HFD (D12331; Research Diets). All experiments were approved by the Danish Animal Experimentation Inspectorate (2018-15-0201-01457). The GDF15 KO and WT littermate mice were generated as described previously (Hsiao et al., 2000). After weaning, mice were housed in groups of two GDF15 KO or WT mice per cage, respectively. Mice were housed at 22±1°C with a 12:12 hour light-dark cycle and ad libitum access to water and chow diet or HFD (Ssniff, E15772-34), respectively. All experiments were approved by the LAVG Brandenburg, Germany (2347-14-2021).

### Pharmacological compounds

Metformin (Sigma, PHR1084) was dissolved in tap water and administered perorally at indicated concentrations using a steel gavage needle in a volume of 5 mL/kg body weight.

### Mouse study 1: Effect of metformin on voluntary running in chow-fed mice

Female WT (n=10) and GFRAL KO (n=10) mice (28-29 weeks of age) were single housed five days prior to experiment in cages equipped with running wheels (23 cm in diameter, Tecniplast, Italy). Running distance was measured by an odometer (Sigma Pure 1 Topline 2016, Germany) and the amount of bedding was reduced in order to avoid blocking the running wheel. On the experimental day, mice were gavaged with vehicle or metformin (400 mg/kg) 3 hours prior the dark phase, and running distance and food intake were measured after 24 hours, followed by a final dose of either vehicle or metformin (400 mg/kg), and plasma for GDF15 analysis was collected after 3 hours. After 11 days of washout with access to the running wheel, a crossover study was conducted, where mice previously gavaged with vehicle received metformin and vice versa. There is one missing value for vehicle-treated GFRAL KO mice for running distance, because of a malfunctioning running wheel (Figure 1e). One food intake value excluded for WT metformin treated group due to shredding (Figure 1d).

### Mouse study 2: HFD-feeding time course

Sixty male mice of the age of 8-10 weeks were housed 3 per cage and fed with standard chow diet. After 1 week of acclimatization, mice were gavaged with either vehicle (n=6) or metformin (400 mg/kg) (n=6). Two hours later, these mice were euthanized by decapitation and blood samples collected on wet-ice for plasma separation and tissues were collected and immediately frozen on dry ice. This procedure was repeated for mice fed HFD for 1 day, 3 days, 10 and 21 days. Plasma samples were analyzed for GDF15 and RNA was extracted from liver, kidney, duodenum and colon and *Gdf15* mRNA expression was measured by qPCR.

### Mouse study 3: *Effect of metformin on voluntary running in HFD-fed mice*

Female WT (n=10) and GFRAL KO (n=10) mice were switched to a HFD following a 16-day washout period after “Mouse Study 1”. Mice had access to running wheels during the washout period. Following 5 days on HFD, mice were gavaged with vehicle or metformin (400 mg/kg) 3 hours prior to the onset of the dark phase and running distance and food intake were measured 24 hours following administration. Next day, 24 hours after the first administration, a final dose of vehicle or metformin, were given and blood samples were obtained 3 hours later. One running distance value is missing for a vehicle treated GFRAL KO mouse, given a malfunctioning running wheel, accordingly the food intake of this animal was also excluded from the analysis (Figure S1D and S1E).

### Mouse study 4: Effect of metformin on HFD-induced weight gain in WT and GFRAL KO mice

Male WT (n=16) and GFRAL KO (n=20) mice (age 11-20 weeks old) were treated p.o. for 11 days with vehicle or 300 mg/kg metformin (3 hours before the onset of the dark phase). Before starting treatment with metformin or vehicle, respectively, mice were fed at HFD and gavaged with tap water daily for 3 days to acclimatize them to handling and p.o. treatment and minimize stress-mediated effects on body weight and food intake during metformin treatment. On day 11, 3 hours following the final administration, plasma was collected for GDF15 assessment. One missing value in GFRAL KO metformin group due to insufficient amount of plasma collected (Figure 2B)

### Mouse study 5: Effect of metformin on HFD-induced weight gain in WT and GDF15 KO mice

Male WT (n=20) and GDF15 KO (n=14) (age 10-16 weeks) mice were treated p.o. for 11 days with vehicle or 300 mg/kg metformin (3 hours before onset of dark phase). Before starting treatment with metformin or vehicle, respectively, mice were gavaged with tap water daily for 3 days to acclimatize them to handling and p.o. treatment and minimize stress-mediated effects on body weight and food intake during metformin treatment. On day 11, 3 hours following the final administration, blood glucose was measured and plasma for GDF15 assessment was obtained.

### Mouse study 6: Effect of metformin on body weight and food intake in WT DIO mice

28 male HFD-fed DIO mice (body weight >50g) were randomly assigned to vehicle (n=7) or metformin (200mg/kg (n=7), 300mg/kg (n=7) or 400mg/kg (n=7)) for 7 days (p.o., 3 hours before onset of dark phase). Before starting treatment with metformin or vehicle, mice were gavaged with tap water daily for 3 days to acclimatize them to handling and p.o. treatment and minimize stress-mediated effects on body weight and food intake during metformin treatment. On day 7, a final dose was delivered and 3 hours later, blood glucose was measured and blood for GDF15 assessment was obtained. Two animals (one from 300mg/kg metformin group and another from 400mg/kg metformin group) were removed from the study due to severe adverse responses to the treatment (constipation, patent apathy). One outlier was detected by Grubbs’ test for GDF15 plasma analysis (200mg/kg group) and one plasma sample was missing for the vehicle group.

### Mouse study 7: Effect of metformin on body weight and food intake in DIO WT and GFRAL KO mice

Male HFD-fed DIO WT (n=10) and GFRAL KO (n=11) mice at the age of 26-27 weeks were treated p.o. for 7 days with vehicle or 400 mg/kg metformin (3 hours before onset of dark phase). Mice were single housed and before starting treatment with metformin or vehicle, respectively, mice were gavaged with tap water daily for 3 days to acclimatize them to handling and p.o. treatment and minimize stress-mediated effects on body weight and food intake during metformin treatment. On day 7, a final dose was delivered and 3 hours later, plasma for GDF15 assessment was obtained.

### Mouse study 8: Effect of metformin on body weight and food intake in DIO WT and GDF15 KO mice

Male HFD-fed DIO WT (n=20) and GDF15 KO (n=14) mice at the age of 24-28 weeks were housed in groups of 1-2 mice/cage and treated p.o. for 7 days with vehicle or 400 mg/kg metformin (3 hours before onset of dark phase). Before starting treatment with metformin or vehicle mice were gavaged with tap water daily for three days to acclimatize them to handling and p.o. treatment and minimize stress-mediated effects on body weight and food intake during metformin treatment. On day 7, a final dose was delivered and 3 hours later, blood glucose was measured and plasma for GDF15 assessment was obtained

### Mouse study 9: Acute effect of metformin on body weight and food intake in DIO WT and GFRAL KO mice

Male HFD-fed single housed DIO WT (n=8) and GFRAL KO (n=11) mice were treated once with vehicle or 400 mg/kg metformin (3 hours before onset of dark phase) and 24 hours after the single administration body weight and food intake were measured. The animals were allowed 5 days to recover before treatment groups were switched (cross-over design) and the experiment was repeated. After 8 additional days of recovery, mice were randomly assigned to receive either vehicle or 400 mg/kg metformin, 3 hours later plasma was obtained for measurement of GDF15 (Figure S2C-F).

### Mouse study 10: Effect of morning-administered metformin on body weight and food intake in DIO WT and GDF15 KO mice

To delineate circadian effects of metformin on food intake and body weight, male mice at the age of 25-29 weeks were treated one hour after the beginning of the light phase (5:30 am). HFD-fed DIO WT (n=20) and GDF15 KO (n=20) mice were housed in groups of two mice/cage and treated p.o. for 7 days with vehicle or 400 mg/kg metformin. Before starting treatment with metformin or vehicle, respectively, mice were gavaged with tap water daily for three days to acclimatize them to handling and p.o. treatment and minimize stress-mediated effects on body weight and food intake during metformin treatment. On day 7, a final dose was delivered and 3 hours later, blood glucose was measured and plasma for GDF15 assessment was obtained (Figure S2G-K).

### GDF15 ELISA

GDF15 was measured in human plasma or serum using the Quantikine ELISA Human GDF-15 Immunoassay (ELISA, R&D systems, catalog no. DGD150). In mice, plasma samples were analyzed using Quantikine ELISA Mouse GDF15 Immunoassay (ELISA, R&D systems, catalog no. MGD150). The ELISA assays were used according to the protocol provided by the manufacturer.

### RNA extraction & cDNA synthesis

Tissue was quickly dissected and frozen on either dry ice or liquid nitrogen and stored at -80°C. Tissue was homogenized in Trizol reagent (QIAzol Lysis Reagent, Qiagen) using a stainless steel bead (Qiagen) and a TissueLyser LT (Qiagen) for 3 min at 20 Hz. Then, 200 μl chloroform (Sigma-Aldrich) was added and tubes were shaken vigorously for 15 sec and left at RT for 2 min, followed by centrifugation at 4°C for 15 min at 12,000 x g. The aqueous phase was mixed 1:1 with 70% ethanol and further processed using RNeasy Lipid Mini Kit following the instructions provided by the manufacturer. For muscle tissue, the lysis procedure, described in the enclosed protocol of the Fibrous Tissue Mini Kit (Qiagen), was followed. After RNA extraction, RNA content was measured using a NanoDrop 2000 (Thermo Fisher) and 500 ng of RNA was converted into cDNA by mixing FS buffer and DTT (Thermo Fisher) with Random Primers (Sigma-Aldrich) and incubated for 3 min at 70°C followed by addition of dNTPs, RNase out, Superscript III (Thermo Fisher) and placed in a thermal cycler for 5 min at 25°C, 60 min at 50°C, 15 min at 70°C, and kept at -20°C until further processing.

### qPCR

SYBR green qPCR was performed using Precision plus qPCR Mastermix containing SYBR green (Primer Design, #PrecisionPLUS). Primers used were Gdf15: F: CCGAGAGGACTCGAACTCAG; R: ACCCCAATCTCACCTCTGGA, Rpl13a: F: GGAGGGGCAGGTTCTGGTAT; R: TGTTGATGCCTTCACAGCGT. qPCR was performed in 384-well plates on a Light Cycler 480 Real-Time PCR machine using 2 min preincubation at 95°C followed by 45 cycles of 60 sec at 60°C and melting curves were performed by stepwise increasing the temperature from 60°C to 95°C. Quantification of mRNA expression was performed according to the delta-delta Ct method.

### Statistical analyses

Statistical analyses were performed in Graphpad Prism (version 9) and Sigma Plot (version 14). For comparing multiple groups, one-or two-way ANOVA or two-way RM ANOVA was used. When ANOVA revealed a significant interaction, Bonferroni post hoc multiple comparisons test was used. When comparing two groups, an unpaired two-tailed Students t-test was used. Data were evaluated for Gaussian distribution and equal variance by Shapiro-Wilk test, Brown-Forsythe test, and by visual inspection of the distribution of residuals. When data failed to follow Gaussian distribution or equal variance after transformation (in the following order: ln(x), log10(x), √(x)), data were analyzed by Mann-Whitney test, Welch’s test, or Kruskal-Wallis on Ranks. Grubbs’ test was employed to determine significant outliers (alpha = 0.05). The analysis of changes in plasma GDF15 for human study 1 was performed using a baseline constrained repeated measures regression model with plasma GDF15 as a function of age, sex, time and a group-by-time interaction. All participants were assigned to the control group at baseline thereby ascribing all between participants differences at baseline to the individual level. Degrees of freedom were computed by the method of Kenward and Rogers and an unstructured covariance was specified. GDF15 data were transformed using the natural logarithm due to the residuals of the regression model being right skewed. Missing data were assumed to be missing at random, or completely at random, in relation to the outcome. This analysis was performed using SAS V.9.4 M (SAS Institute). Unless otherwise stated, all data are presented as mean ± SEM. P < 0.05 was considered statistically significant.

