## Supplementary material for "The GDF15-GFRAL pathway is dispensable for the effects of metformin on energy balance": Combined Supplemental Data

**Figure S1**

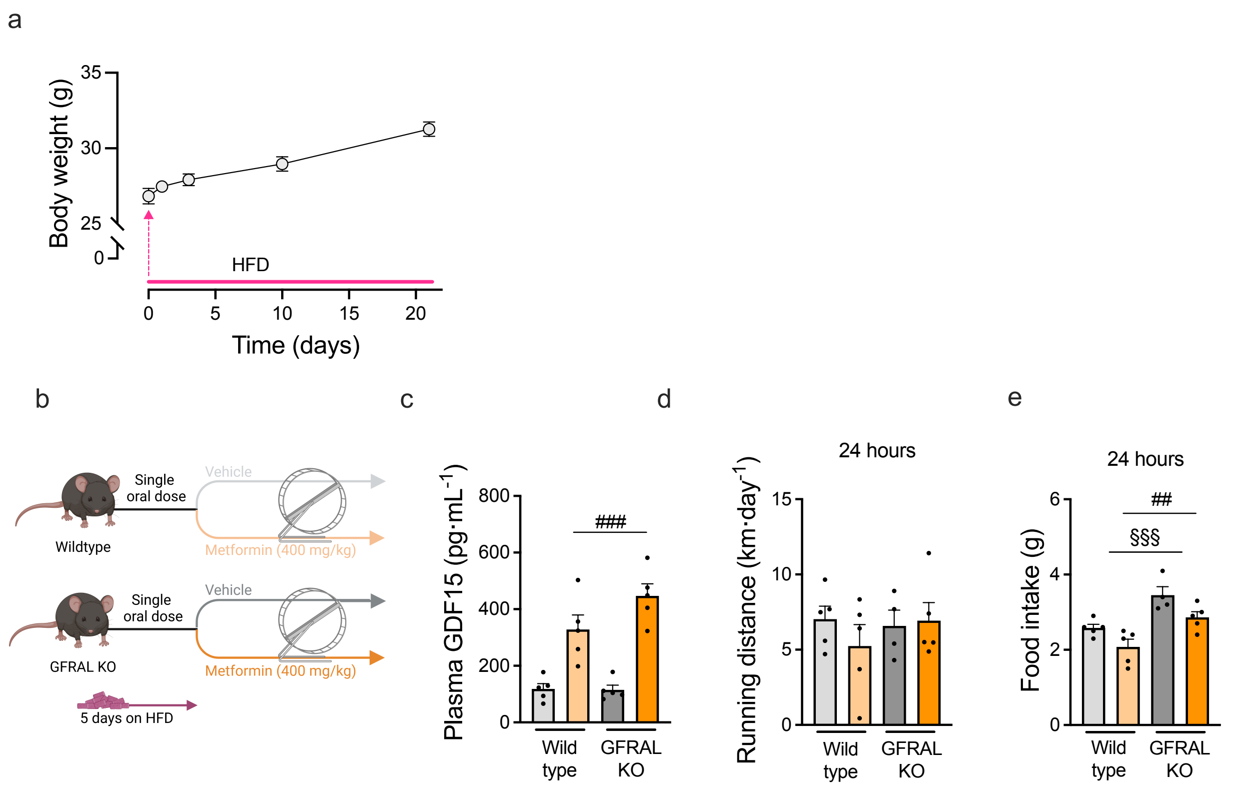


**Figure S1. Metformin increases circulating GDF15 and lowers food intake but does not affect running distance in female mice. a,** Body weight gain of mice used for high-fat diet (HFD) time-course with metformin. **b**, Illustration of study in which female lean wild type and GFRAL knockout (KO) mice were fed HFD for 5 days while having access to running wheels. Mice were treated with metformin (400 mg/kg) or vehicle by oral gavage and running distance over 24 hours was measured. **c,** Blood plasma GDF15 levels 3 hours following metformin (400 mg/kg). Data shown as mean ± s.e.m and analyzed by a two-way ANOVA, $^{###}p<0.001$; main effect of metformin. **d**, Running distance after 24 hours in mice from b. Data shown as mean ± s.e.m and analyzed by a two-way ANOVA. **e,** Food intake after 24 hours. Data shown as mean ± s.e.m and analyzed by a two-way ANOVA, $^{§§§}p<0.001$; main effect of genotype, $^{##}p<0.01$; main effect of metformin.

**Figure S2**

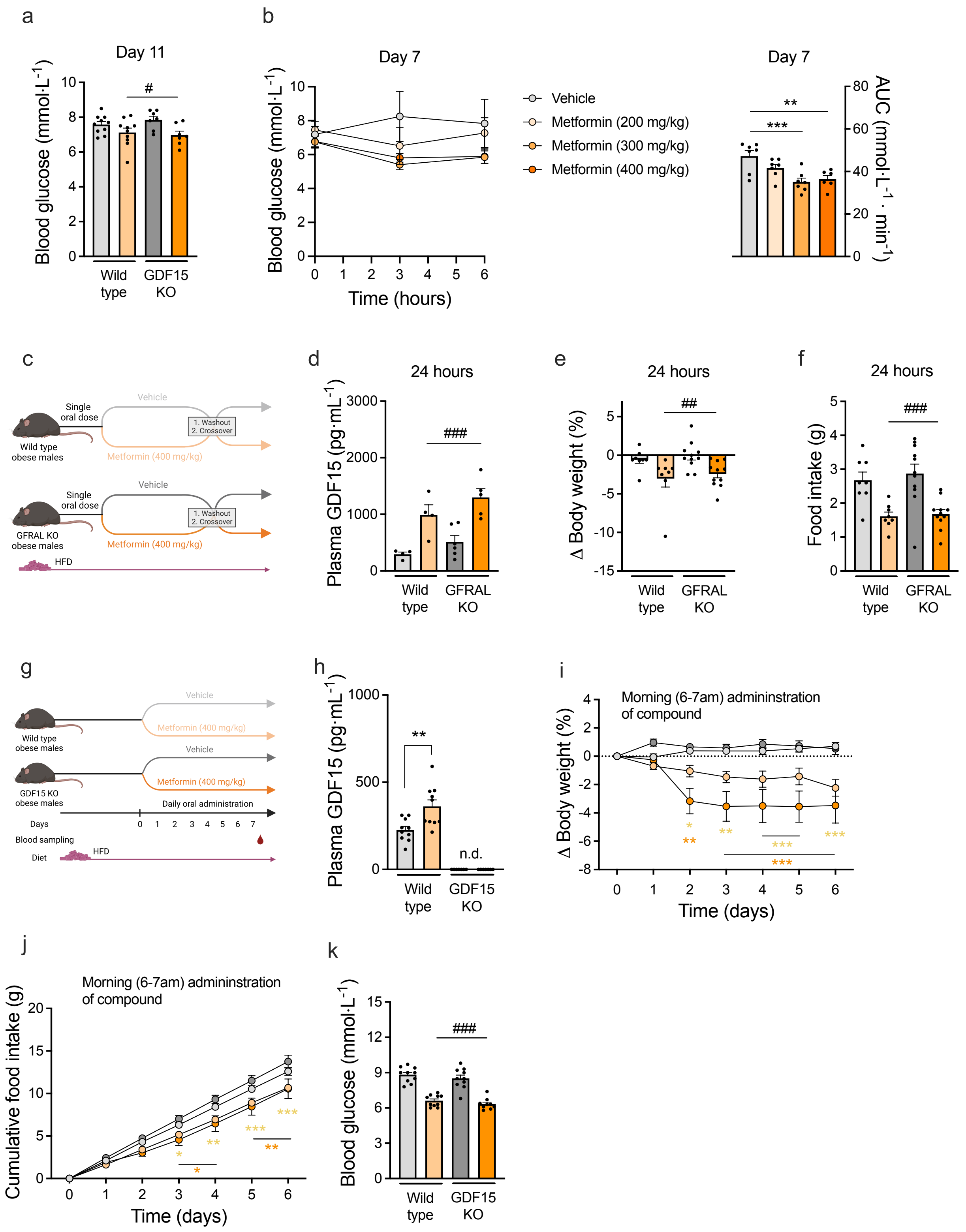
a
Day 11
Blood glucose (mmol·L-1)
#
Wild type
GDF15 KO
b
Day 7
Blood glucose (mmol·L-1)
Time (hours)
Vehicle
Metformin (200 mg/kg)
Metformin (300 mg/kg)
Metformin (400 mg/kg)
Day 7
AUC (mmol·L-1 · min-1)
**
***
c
Single oral dose
Vehicle
1. Washout 2. Crossover
Wild type obese males
Metformin (400 mg/kg)
GFRAL KO obese males
HFD
d
24 hours
Plasma GDF15 (pg·mL-1)
###
Wild type
GFRAL KO
e
24 hours
Δ Body weight (%)
##
f
24 hours
Food intake (g)
###
g
Wild type obese males
GDF15 KO obese males
Daily oral administration
Days
Blood sampling
Diet
h
Plasma GDF15 (pg·mL-1)
**
n.d.
i
Morning (6-7am) admininstration of compound
Δ Body weight (%)
Time (days)
j
Cumulative food intake (g)
k
Blood glucose (mmol·L-1)

**Figure S2. Metformin improves glycemic control and lowers body weight independently of the GDF15-GFRAL pathway. a,** Male wild type and GDF15 knockout (KO mice were switched from chow diet to high-fat diet (HFD) and following 3 days on HFD, metformin (300 mg/kg) was orally administered daily for 11 days. On the last day (day 11), blood glucose was determined 3 hours following the final dose of metformin. Data shown as mean ± s.e.m and analyzed by a two-way ANOVA, $^{\#}p<0.05$; main effect of metformin. **b,** Daily oral administration of vehicle or metformin (200, 300, or 400 mg/kg) to male high-fat diet-induced obese mice for 6 days. The next day (day 7) blood glucose was assessed following which another dose of vehicle or metformin (200, 300, or 400 mg/kg) was administered and blood glucose was measured 3 and 6 hours after administration. Right side displays Area Under Curve (AUC) of left side. Data shown as mean ± s.e.m and analyzed by a one-way ANOVA with Bonferroni multiple comparison test (AUC), $^{**}p<0.01$, $^{***}p<0.001$. **c**, Illustration of study in which high-fat diet (HFD)-induced obese wild type and GFRAL knockout (KO) mice received a single dose of vehicle or metformin (400 mg/kg) by oral gavage followed by a 16-day washout period and cross-over of treatments. **d**, Blood plasma GDF15 levels 3 hours after a final administration of vehicle or metformin (400 mg/kg). Data shown as mean ± s.e.m and analyzed by a two-way ANOVA, $^{\#\#\#}p<0.001$; main effect of metformin. **e**, Change in body weight. Data shown as mean ± s.e.m and analyzed by a two-way RM ANOVA, $^{\#\#}p<0.01$; main effect of metformin. **f**, Food intake. Data shown as mean ± s.e.m and analyzed by a two-way RM ANOVA, $^{\#\#\#}p<0.001$; main effect of metformin. **g**, Illustration of study in which high-fat diet (HFD)-induced obese male wild type and GDF15 knockout (KO) mice for seven days received daily administration of vehicle or metformin (400 mg/kg) by oral gavage in the morning at 6.00 am. **h**, Blood plasma GDF15 levels 3 hours after the final dose of vehicle or metformin on day 7. The GDF15 plasma levels were below the detection limit for GDF15 KO mice (n.d. = not detectable). Data shown as mean ± s.e.m and analyzed by a Welch's test within the wild type group, $^{**}p<0.01$. **i,** Change in body weight. Data shown as mean ± s.e.m and analyzed by a two-way RM ANOVA with Bonferroni multiple comparison test, $^{*}p<0.05$, $^{**}p<0.01$; $^{***}p<0.001$; effect of metformin within genotype. **j,** Cumulative food intake. Data shown as mean ± s.e.m and analyzed by a two-way RM ANOVA with Bonferroni multiple comparison test, $^{*}p<0.05$, $^{**}p<0.01$; $^{***}p<0.001$; effect of metformin within genotype. **k,** Effect on blood glucose on day 7, 3 hours after the final drug administration. Data shown as mean ± s.e.m and analyzed by a two-way ANOVA, $^{\#\#\#}p<0.001$; main effect of metformin.

**Figure S3**

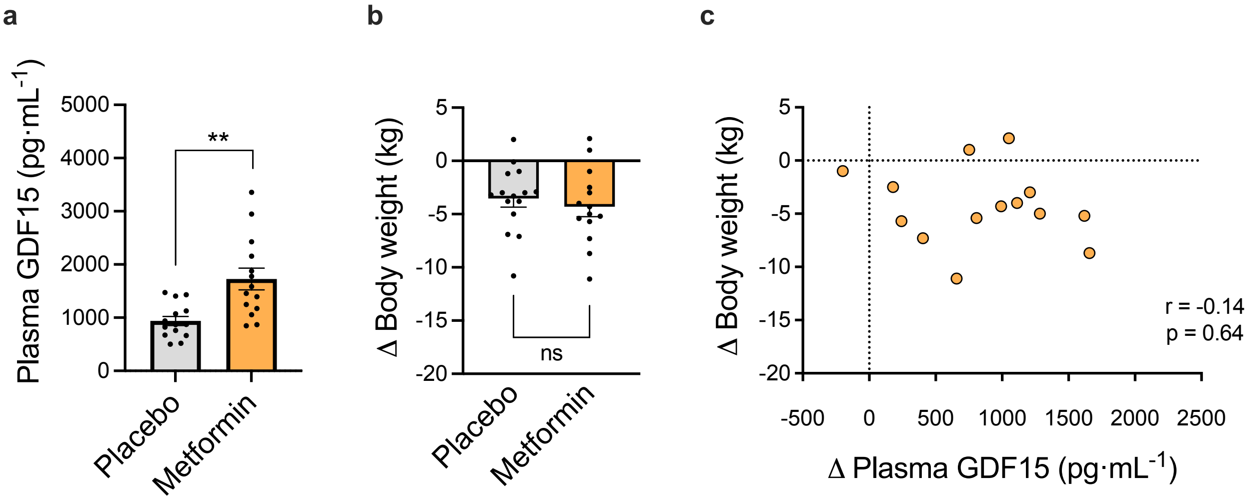


**Figure S3. Metformin in combination with exercise increases circulating GDF15 but is not associated with body weight loss in subjects with overweight/obesity and prediabetes. a,** Blood plasma levels of GDF15 in human subjects with overweight/obesity and prediabetes before and after 12 weeks of exercise training and treatment with either placebo or metformin (1000 mg, twice daily). Data shown as mean ± s.e.m and analyzed by a by a Welch's test between placebo and metformin, **p<0.01. **b,** Change in body weight following 12 weeks of exercise training and treatment with placebo or metformin (1000 mg, twice daily). Data shown as mean ± s.e.m and analyzed by an unpaired two-tailed t-test between placebo and metformin, ns=not significant. **c,** Change in body weight versus change in blood plasma GDF15 levels in the metformin-treated group. Data shown as values for each human subject and analyzed by a Spearman correlation test.
